# Stability of representational geometry across a wide range of fMRI activity levels

**DOI:** 10.1101/266585

**Authors:** Spencer A. Arbuckle, Atsushi Yokoi, J. Andrew Pruszynski, Jörn Diedrichsen

## Abstract

Fine-grained activity patterns, as measured with functional magnetic resonance imaging (fMRI), are thought to reflect underlying neural representations. Multivariate analysis techniques, such as representational similarity analysis (RSA), can be used to test models of brain representation by quantifying the representational geometry (the collection of pair-wise dissimilarities between activity patterns). One important caveat, however, is that non-linearities in the coupling between neural activity and the fMRI signal may lead to significant distortions in the representational geometry estimated from fMRI activity patterns. Here we tested the stability of representational dissimilarity measures in primary sensory-motor (S1 and M1) and early visual regions (V1/V2) across a large range of activation levels. Subjects were visually cued with different letters to perform single finger presses with one of the 5 fingers at a rate of 0.3-2.6 Hz. For each stimulation frequency, we quantified the difference between the 5 activity patterns in M1, S1, and V1/V2. We found that the representational geometry remained stable, even though the average activity increased over a large dynamic range. These results indicate that the representational geometry of fMRI activity patterns can be reliably assessed, largely independent of the average activity in the region. This has important methodological implications for RSA and other multivariate analysis approaches that use the representational geometry to make inferences about brain representations.

## 1. Introduction

Multivariate analysis of activity patterns has profoundly changed functional magnetic resonance imaging (fMRI) data analysis. Traditional fMRI studies have examined differences in overall activity levels in extended brain regions. In this approach, local fine-grained patterns of activity are removed by smoothing, as they are typically not consistent across individuals. However, it was realized that one could decode the experimental condition from activity patterns within individuals, even if the average activity is the same (Haxby et al., 2001). Decodability can be taken as evidence that the region represents something about the underlying distinction between conditions (Haxby et al., 2014). More importantly, the degree to which different pairs of activity patterns are dissimilar tells us about the structure of the representation. For example, a region involved in the identification of object categories should show large dissimilarities between activity patterns associated with objects from different categories, but smaller dissimilarities between different objects of the same category (Kriegeskorte et al., 2008). The relationship between all pair-wise dissimilarities defines what we call the representational geometry. Representational similarity analysis (RSA, Kriegeskorte et al. 2008), pattern component modelling (PCM, Diedrichsen et al. 2017), and encoding models (Naselaris et al., 2011) all analyze this representational geometry to test between models of brain representations (Diedrichsen and Kriegeskorte, 2017).

When testing representational models with fMRI data, we base our analysis on the Blood Oxygenation Level Dependent (BOLD) signal. To what degree can we make inferences about neural representations using this indirect measure of neural activity? There are a number of reasons why representational fMRI analysis may be limited (see discussion). One important problem, the main focus of this paper, is that the measured representational geometry may depend strongly on the overall activity in a region. This is of concern as we often make comparisons across regions, participants, or attentional states with different activity levels. Although RSA per se is independent of average activity, non-linearities between neural activity and BOLD signal may distort the measured representational geometry. The relationship between the pair-wise pattern dissimilarities will only remain the same if the voxel activities in a region all scale by the same constant as neural activity increases. However, voxels that already have high activity may not scale in the same manner as voxels with low activity. For example, voxels with high activity may saturate. This phenomenon would be comparable to the disappearing details in bright areas of an over-exposed photograph, and could profoundly change the measured representational geometry.

In addition, changes in the spatial point-spread function of the fMRI signal (O’Herron et al., 2016) with increasing activity could have a distorting influence. These non-linear effects would make it difficult to make inferences about the representational content of neural population codes using fMRI data. If these distortions are severe, it would constitute a major problem for representational analyses in general. Therefore, it is important to empirically test if representational geometry remains stable across a wide range of overall activity levels.

We investigated this question with an experiment that allowed us to assess simultaneous patterns in sensory-motor regions (S1 and M1) and in primary and secondary visual cortices (V1 and V2) using RSA. Ejaz et al. (2015) demonstrated a stable representational geometry across humans for individual finger movements in M1 and S1, in which the thumb had the most distinct activity pattern and neighboring fingers showed higher similarities than non-neighboring fingers. Similarly, it has been shown that different complex shapes, such as letters, can be decoded from activity in visual cortices (Miyawaki et al., 2008), as well as colors (Brouwer and Heeger, 2009). We therefore cued finger presses with colored letter cues presented on a screen. We chose a specific letter and color for each finger, such that the representational geometry of perceptual similarities between letters and colors would be different from those in motor regions. To increase the overall activity in both regions, we increased the letter flashing and finger pressing frequency from 0.3Hz to 2.6Hz. For individual finger presses on our isometric device (see section 2.2), a rate of 2.6Hz is close to the upper performance limit.

In interpreting the results, it is important to distinguish between two types of non-linearities. The first is the non-linearity between behavior (pressing) or stimuli (flashed digits) and neural activity. This non-linearity is of no concern here – rather it is a feature of the system that we would like to understand using representational models. The second type of non-linearity may exist between neural activity and the BOLD signal. This is the non-linearity we are concerned about. While some studies suggest that this relationship is fairly linear in M1/S1 (Siero et al., 2013) and V1 (Boynton et al., 1996; Heeger et al., 2000), these studies have only looked at the overall activity level. For representational analyses, however, it is important that this holds true for individual voxels, such that the representational geometry remains the same.

## 2. Methods

### 2.1. Participants

We measured cortical activity patterns in 5 female and 3 male participants (mean Age = 25.5(2.41) years). All participants were self-reported right handers (mean Edinburgh questionnaire laterality quotient = 91.25(7.82)), and made individuated finger presses of the right hand.

### 2.2. Apparatus and stimuli

The motor behavior was monitored by a keyboard-like device. The device had a key for each finger of the right hand, with a force transducer (Honeywell-FS series, dynamic range = 0-16N, resolution <0.02N) mounted under each key. A bevel on each key ensured fingertip placement across participants was consistent. Forces were recorded at a sampling rate of 200Hz.

To evoke visual responses, we presented colored letter stimuli on the screen at the same frequency as the finger presses. The aim was to pick letters and colors that would induce a dissimilarity structure that would differ considerably from that of the fingers. Therefore, we chose similar letters and colors for fingers which evoke different activity profiles (see Fig. 1a for finger-letter-color pairs). The letters were presented centrally and peripherally (see Fig. 1b). The size of the letters on the screen were 8 x 10 cm, subtending a visual angle of approximately 70°. The screen background was black. Participants were instructed to maintain visual fixation on a grey cross presented centrally on the screen. Cues were presented centered on the fixation cue. Five lines were presented in the lower third of the screen, one for each of the five fingers. The locations of these lines were dynamically updated to indicate the real-time force applied to each key of the fingerboard device.

**Figure 1:**
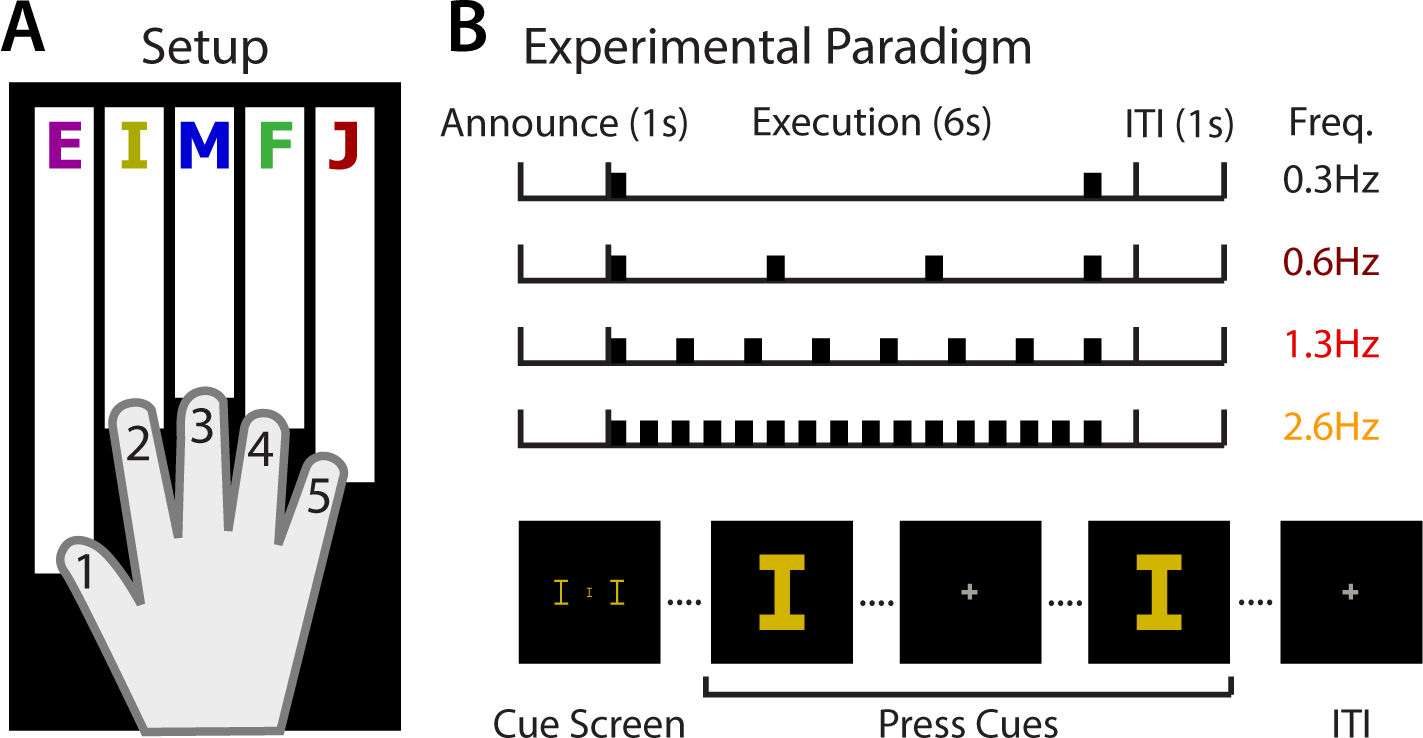
Experimental paradigm. (A) Participants made short, isometric presses of an individual finger onto a keyboard while in an MR scanner. Each finger press was cued with a unique color-letter combination. (B) A cue at the start of each trial (1s) instructed participants which finger they would press. Participants then executed presses when prompted by a larger cue presentation. The cues flashed either 2, 4, 8, or 16 presses in 6 seconds (0.3, 0.6, 1.3, 2.6Hz). A 1s inter-trial-interval (ITI) separated each trial. Random periods of rest were interleaved between trials in each block. This design yielded 20 conditions (5 fingers/letters x 4 frequencies).

### 2.3. Behavioral task

In the MR scanner, participants completed a paced finger pressing task. Each trial lasted for 8 seconds and was divided into three phases (see Fig. 1b). In the announce phase (1 second), participants saw a visual cue indicating which finger of their right hand they were to press in the current trial. During the following execution phase (6 seconds), participants made 2, 4, 8, or 16 isometric presses of the finger (0.3, 0.6, 1.2, and 2.3Hz pressing frequency). Each press was paced by a visual metronome, which flashed the letter cue from the announce phase for 100ms each. The first and the last press always occurred at the beginning and end of the execution phase, with the intermediate presses being cued at a constant rate (see Fig. 1b). Following this, there was an inter-trial interval of 1 second before the next trial started. Throughout the entire experiment, participants were instructed to refrain from moving their wrist or fingers of either hand when not instructed to do so.

There were 20 possible conditions (5 fingers/letters x 4 pressing frequencies). The task was divided into 8 runs of 40 trials each, with two repeats per condition. Trial order within each run was randomized. Seven periods of rest (13 seconds) were randomly interspersed between trials in each run. Each run lasted 411 seconds. Participants were instructed to produce a minimum of 2N, but not to exceed a maximum of 4N with each finger press. Minimum and maximum force thresholds were visually presented on screen above the finger force lines. Between presses, the force applied to each key needed to be below 0.75N before another press could be registered. The fixation cross turned white for each correct press. Participants trained on this task prior to the scanning portion of the experiment to ensure stable performance.

### 2.4. fMRI Data Acquisition

Functional images were acquired using a Siemens Magnetom Syngo 7T MRI scanner with a 32-channel head coil at the University of Western Ontario (London, Ontario, Canada). Volumes were acquired using an interleaved, multi-band slice acquisition (TR = 1000ms, 44 slices, 1.4mm isotropic voxels, no gap between slices, in-plane acceleration factor = 3, multi-band factor = 4). The first three images of each functional run were discarded to allow magnetization to reach equilibrium. The slices covered the dorsal aspects of the cerebrum, encompassing M1 through to V1. A T1-weighted anatomical scan (3D MPRAGE sequence, TR = 6000ms, 0.75mm isotropic voxels, 208 volumes) was also acquired at the start of the scan. Fieldmaps were collected at the end of the imaging session.

### 2.5. Preprocessing and first-level model

First-level fMRI analyses were conducted with SPM12 (http://www.fil.ion.ucl.ac.uk/spm/software/spm12/). Functional images were realigned to correct for motion across runs. Within this process, we utilized a B0 fieldmap to correct for magnetic field inhomogeneities. Due to the short TR, no slice timing corrections were applied. The functional data was co-registered to the individual anatomical scan, but no normalization was applied.

The pre-processed images were analyzed with a general linear model (GLM) with separate task regressors for each condition (20 regressors) for each run. Each regressor was a boxcar function that was on for 6 seconds of the trial duration and off otherwise. These regressors were then convolved with a hemodynamic response function with a peak onset of 4.5 seconds and a post-stimulus undershoot minimum at 11 seconds.

We used the SPM FAST autocorrelation model in conjunction with restricted-maximum likelihood (ReML) estimation to estimate the long-range temporal dependencies in the functional timeseries. The coefficients of this model were estimated from all grey-matter voxels. We then used the temporal covariance estimate to temporally prewhiten the time-series data. Because this step effectively attenuates low-spatial frequencies, it removes the necessity to apply a separate high-pass filter. We found that on a number of data sets, this analysis procedure improves the reliability of activity pattern estimates as compared to the standard high-pass filtering and subsequent temporal autocorrelation correction with FAST.

### 2.6. ROI definitions

We used Freesurfer software (Dale et al., 1999) to extract the white-gray matter and pial surfaces from each participant’s anatomical image. These surfaces were inflated to a sphere and aligned using sulcal depth and curvature information to the Freesurfer average atlas (fsaverage, Fischl et al. 1999). Following alignment, both hemispheres in each participant were resampled into a 163,842 vertex grid. This allowed us to reference similar areas of the cortical surface in each participant by selecting the corresponding vertex on the group atlas.

Anatomical regions of interest (ROI) were defined using a procedure established in previous work (Wiestler and Diedrichsen, 2013; Ejaz et al., 2015). All ROIs were defined using a probabilistic cytoarchtectonic atlas (Fischl et al., 2008) projected onto the common group surface. For M1 and S1, we constrained the resulting ROIs to the hand and arm region by choosing the area of the cyctoarchteconically defined strip 2cm above and below the hand knob (Yousry et al., 1997). To avoid cross-contamination between M1 and S1 activities along the central sulcus, voxels with more than 25% of their volume originating from the opposite side of the central sulcus were excluded. The primary visual cortices (V1/V2) were grouped as one ROI. The group area was then projected onto the individual volume using the individual surface reconstruction.

### 2.7. Multivariate fMRI analysis

Multi-voxel analyses were conducted within each ROI (M1, S1, and V1/V2), using the RSA (Nili et al., 2014) and PCM toolboxes (Diedrichsen et al., 2017). For each ROI, we extracted the beta-weights from the first-level GLM for each condition in each imaging run. These beta-weights were then spatially pre-whitened using multivariate noise-normalization to suppress correlated noise across voxels (Walther et al., 2016). The mean pattern was not removed from each run to preserve information about activity relative to baseline in each voxel.

We then calculated the squared cross-validated Mahalanobis distance (crossnobis; Walther et al. 2016; Diedrichsen et al. 2016) between activity patterns:

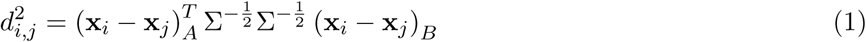

where (*x_i_* − *x_j_*)_*A*_ correspond to the difference between the activity patterns of condition *i* and *j* in run *A*. We repeated this procedure over all possible leave-one-run-out crossvalidation folds and then averaged the resulting dissimilarities across folds. This procedure leads to an unbiased distance estimate, which on average will be zero if there is no reliable difference between the two patterns. This also means that the crossnobis estimator can become negative. The large advantage for this measure, however, is that zero is meaningfully defined and hence ratios between distances can be interpreted in a meaningful way.

The representational geometry is characterized by the dissimilarities between all possible pairs of condition activity patterns, that can be collected into a representational dissimilarity matrix (RDM). The RDM is a (number of conditions x number of conditions) symmetric matrix, with zeros along the diagonal. Dissimilarities were calculated for the left hemisphere M1 and S1 ROIs (contralateral to the side of finger movements). For V1/V2 ROIs, we first calculated dissimilarities for each hemisphere, then averaged the dissimilarities across hemispheres within each participant. We used classical multi-dimensional scaling (eigen-value decomposition) to visualize a low-dimensional projection of the representational geometry (Diedrichsen et al., 2017).

### 2.8. Stability of the representational geometry across stimulation-frequencies

To assess the stability of the representational geometry across frequency conditions, we correlated the RDMs across all 6 possible frequencies pairs within each participant. For each frequency condition (*j*), we had 10 pairwise dissimilarities (*d_i,j_*). We calculated a Pearson correlation without subtracting the mean across the 10 distances first, as zero is a fixed and meaningful value for the unbiased crossnobis distance (see above). The RDM correlation between condition *j* and *k* then becomes

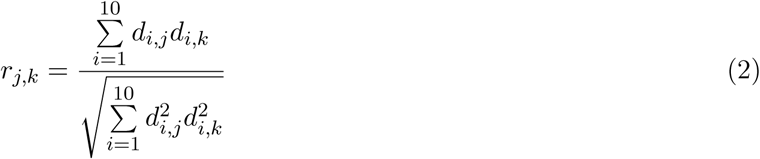

Because the RDM is a symmetric matrix, we correlated only the dissimilarities in the lower triangular of each RDM (excluding the zero diagonal values). To compare the correlation values to a meaningful noise ceiling (see section 2.9), we split the data into odd and even runs and calculated crossnobis distances for each partition separately. We then calculated the correlation either between odd runs for condition *j* and even runs for condition *k*, or the other way around. The two correlations were then averaged for each participant and cross-frequency pair.

### 2.9. Reliability of representational geometries

Even if the representational geometry was perfectly stable across frequencies, the resultant correlations would not be 1 given the noise in our measurements. Therefore, to interpret these cross-frequency RDM correlations meaningfully, we used the reliability of each RDM to estimate a noise-ceiling for each of the 6-possible cross-frequency pairs. We measured the split-half reliability of the RDM at each frequency (*r_j_*, *r_k_*) using the same procedure used to calculate the cross-frequency correlations, but this time correlating the RDMs for odd and even runs within frequencies. If the activity patterns of condition j and k are identical, the expected cross-frequency correlation would be

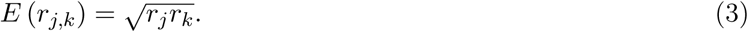

This prediction provides an appropriate noise ceiling for the measured cross-frequency correlations. We calculated the difference between the measured cross-frequency correlations and the corresponding noise ceilings, and tested the deviations using a one-tailed sign-test. Deviations significantly lower than zero indicated that the correlations between the cross-frequency RDMs were lower than expected given the reliability of each RDM.

## 3. Results

We measured cortical BOLD activity patterns as participants saw digits flashed repeatedly on the screen (Fig. 1b), and made short, isometric presses of each finger of the right hand (Fig. 1a). We systematically increased the stimulation and pressing frequencies (Fig. 1b) to increase the overall activity in the visual and motor regions. Our main question was whether the representational geometry – i.e. the collection of relative dissimilarities between different conditions, would remain relatively stable across a large range of overall activity. Behaviorally, participants were able to follow the pacing and to perform the task with relatively high accuracy and matched forces (see Table 1).

Figure 2a shows a surface representation of the activity patterns in the left hemisphere hand region of primary motor and sensory cortices (M1 and S1) from one participant. The overlapping nature of the activity patterns for the different fingers is clearly visible, as well as an overall gradient with the thumb activating more ventral and the little finger more dorsal aspects of the hand region. The overall intensity of the activity increased with increasing pressing frequency, but the spatial distribution of the activity associated with each finger movement appeared to remain stable. Overall, we observed an approximately 5-fold increase in M1 and S1 (Fig. 2b), with less activity evoked in visual cortices (V1/V2).

In M1 and S1 the relationship between pressing frequency and BOLD was visibly not linear. To quantify this observation, we compared a linear fit between the BOLD activity and pressing frequency with a log-linear fit. We compared the proportion of variance explained in each participant between these two fits, after accounting for the proportion of variance explained by only the mean activity. Activity in M1 was better fit by the log-linear model than the linear model in six of eight participants (average log-linear *R*^2^ = 0.916(0.028) vs. 0.663(0.166), *t*_(7)_ = 1.62, *p* = 0.074), and in half of the participants in S1 (average log-linear *R*^2^ = 0.844(0.053) vs. 0.646(0.158), *t*_(7)_ = 1.20, *p* = 0.134). This non-linear relationship can likely be explained by a non-linear mapping between behavior and neural activity, as successive finger presses executed briefly one-after-another elicit less high-gamma activity (Siero et al., 2013). In V1/V2 the linear and log-linear models accounted for very similar proportions of variance (average log-linear *R*^2^ = 0.384(0.307) vs. 0.362(0.432), *t*_(7)_ = 0.15, *p* = 0.441), either because the relationship between stimulation frequency and neural activity is more linear in this region (up to 5 Hz, see Liu et al. 2010), or because the visual activity did not increase to the same level as in M1 and S1.

**Table 1:** Mean and (in parentheses) between-subject standard error of behavioral measure of the finger pressing task. The pressing frequency is reported in Hertz (Hz), and forces in Newtons (N). Participants (*n* = 8) were able to approximately match the instructed frequency and keep the pressing forces relatively stable.

| Cued frequency: | 0.3Hz |  | 0.6Hz |  | 1.3Hz |  | 2.6Hz |  |
| --- | --- | --- | --- | --- | --- | --- | --- | --- |
| Finger | Pressing<br>freq. (Hz) | Force (N) | Pressing<br>freq. (Hz) | Force (N) | Pressing<br>freq. (Hz) | Force (N) | Pressing<br>freq. (Hz) | Force (N) |
| Thumb | 0.33(0.02) | 3.04(0.08) | 0.66(0.03) | 3.12(0.05) | 1.32(0.04) | 3.18(0.07) | 2.65(0.10) | 3.13(0.07) |
| Index | 0.32(0.03) | 2.82(0.07) | 0.65(0.04) | 2.91(0.07) | 1.33(0.02) | 2.99(0.05) | 2.67(0.21) | 2.97(0.07) |
| Middle | 0.34(0.02) | 3.62(0.16) | 0.67(0.02) | 3.53(0.14) | 1.34(0.03) | 3.52(0.14) | 2.70(0.14) | 3.39(0.11) |
| Fourth | 0.33(0.04) | 2.94(0.08) | 0.67(0.04) | 2.99(0.05) | 1.34(0.03) | 3.07(0.05) | 2.63(0.24) | 2.98(0.07) |
| Little | 0.33(0.02) | 2.99(0.06) | 0.65(0.04) | 2.99(0.05) | 1.32(0.05) | 3.11(0.04) | 2.66(0.21) | 2.97(0.05) |

Our main interest, however, was not whether the relationship between behavior and BOLD signal was linear, but whether possible non-linearities between the neural activity and the patterns of BOLD activation would lead to distortions of the representational geometry. To assess this, we calculated the dissimilarity between pairs of activity patterns for each condition of the same frequency (i.e. the representational geometry), for each ROI in each participant. We then examined how stable this geometry remained despite the nearly 5-fold increase in overall activity. As a first step, we visualized the group average representational geometry in M1 and V1/V2 using a multi-dimensional scaling plot.

For M1 (Fig. 3a), we observed the expected representational geometry with the thumb having the most unique pattern and the other fingers being arranged according to their neighborhood relationship (Ejaz et al., 2015). As pressing frequency increased, this arrangement scaled up and substantially moved away from resting baseline (Fig. 3a, cross), but the overall geometry remained the same. This stability can also be appreciated when visualizing the 10 pairwise differences between fingers for each frequency (Fig. 4a,e).

**Figure 2:**
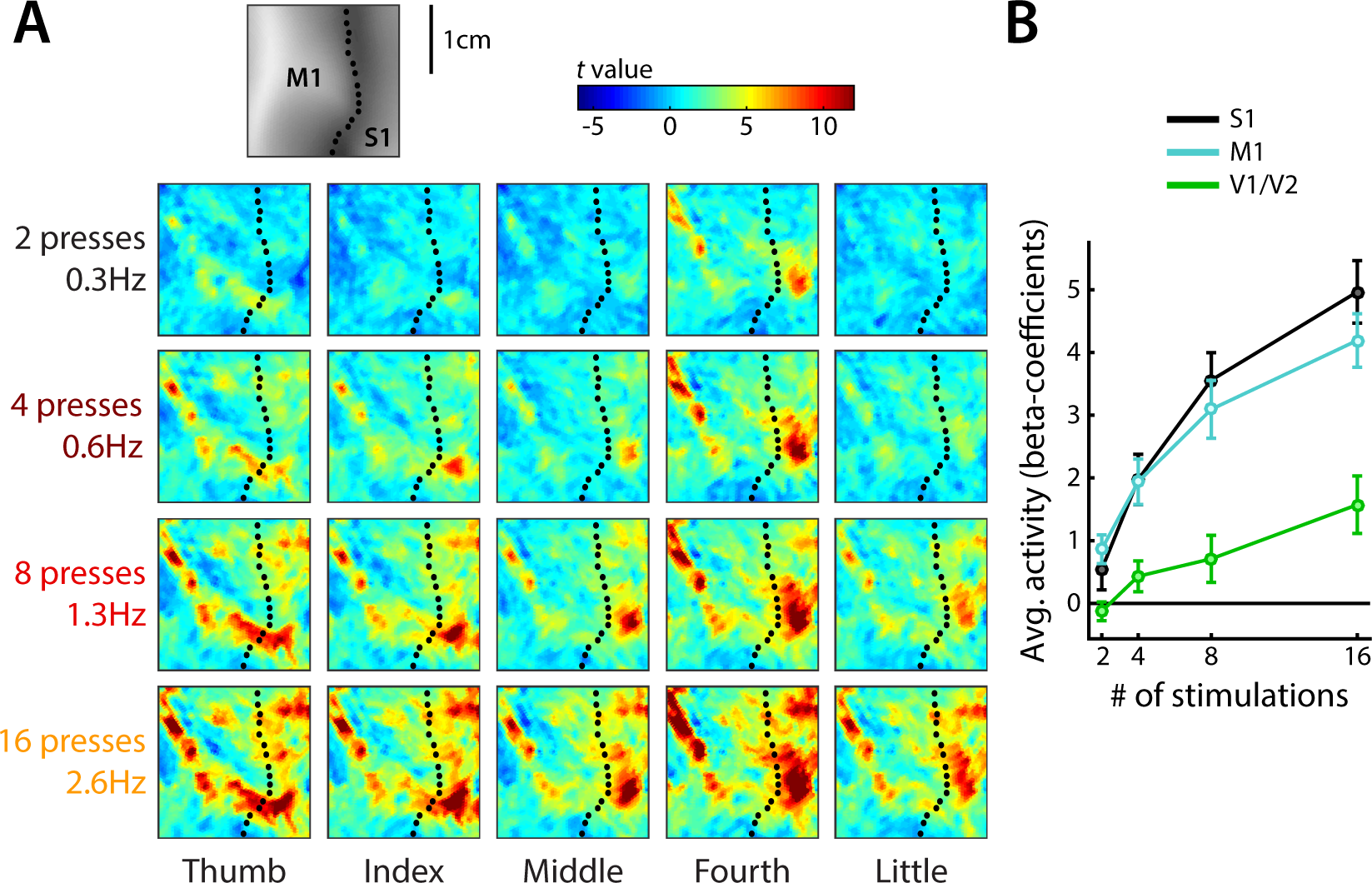
Scaling of activity patterns. (**A**) BOLD activity patterns from the hand area of the primary sensorimotor cortices of an example participant projected onto a flat, surface reconstruction of their cortex. Dotted lines indicate the fundus of the central sulcus. The top insert reflects sulcal depth (darker colors reflect larger depths) and denotes location of M1 and S1. Color maps reflect t-values of activity against rest. Each column corresponds to one finger (thumb to little), and each row one pressing frequency (0.3 to 2.6Hz). The activity increases with increasing pressing frequency. (**B**) Average activity (beta-coefficients) in contralateral M1, S1, and bilateral V1/V2 as a function of pressing/stimulation frequency. Error bars reflect s.e.m.

In the visual cortices (V1/V2), we observed a distinct arrangement of the conditions, with the representational geometry relating to the letters and colors presented for each finger. Similarly to M1 and S1, this structure scaled up with increasing stimulation frequency (Fig. 3b). However, the two lowest stimulation frequencies were not very successful in eliciting either average activity or reliable activity patterns (see below).

To quantify the stability of the representational geometry across different levels of activity, we correlated the dissimilarities in M1, S1, and V1/V2 (Fig. 4a,e,i) across frequencies within each participant. The average cross-frequency correlations (Pearson correlation without intercept- see section 2.8) was *r* = 0.92 in M1 (Fig. 4b) and *r* = 0.94 in S1 (Fig. 4f). In V1/V2 (Fig. 4j), cross-frequency correlations were low when they involved the lowest stimulation frequency, but increased with stimulation frequency (the average cross-frequency correlation between the RDMs of the two highest frequencies was 0.9).

**Figure 3:**
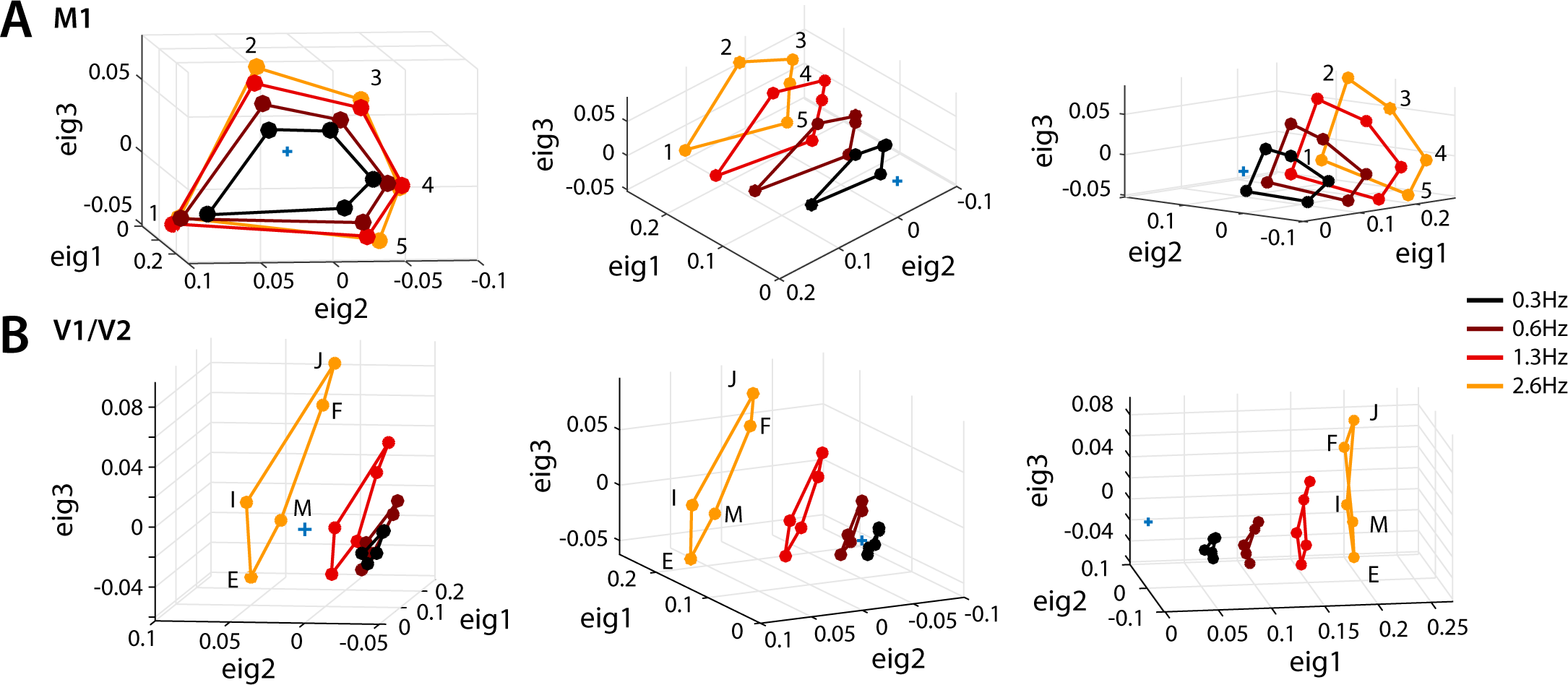
Multidimensional scaling of condition activity patterns for M1 (**A**) and bilateral V1/V2 (**A**). In both M1 (**A**) and V1/V2 (**A**), the patterns scale with increasing pressing/stimulation frequency. The arrangement of these patterns (the representational geometry) remains approximately stable, and expands as the frequency increases. Colors correspond to pressing/stimulation frequency. Numbers indicate fingers (1 = thumb, 5 = little finger). Letters correspond to the flashed letter cue (1=E, 2=I, 3=M, 4=F, 5=J). Note that the conditions are connected differently for M1 and V1/V2. Panels in each row present a different field of view of the same plot. The blue cross indicates baseline.

The cross-frequency correlations are expected to be smaller than 1 even if they are perfectly stable, given the measurement noise in the data. Therefore, to quantify the stability of the representational geometry, we calculated a noise-ceiling (the expected correlation if the true patterns were identical across frequencies- see section 2.9) for each cross-frequency pair. We first determined the split-half reliabilities of the RDMs within each frequency. In M1 and S1, the average split-half reliabilities across participants (Fig. 4c,g) was high (*r >* 0.9) for all pressing frequencies. In V1/V2 (Fig 4k), the reliabilities of the dissimilarities measured at the slowest stimulation frequencies were lower, likely due to low levels of evoked activity, but increased comparably to reliabilities measured for M1 and S1 at higher frequencies. We then estimated the noise-ceilings for each cross-frequency pair by calculating the geometric means of pairs of split-half reliabilities for different frequencies within each participant (see section 2.9).

Comparison between the measured and the expected (noise ceiling) correlations confirmed that the representational geometries remained as stable as could be expected based on the level of measurement noise across a broad range of overall activities. The right-most column in Figure 4 shows the differences between the measured cross-frequency correlations and their respective noise-ceilings. Negative values indicate that the measured correlations were lower than the estimated noise-ceiling for that cross-frequency pair. In V1/V2 (Fig. 4l), one-tailed sign-tests indicated the measured cross-frequency correlations did not significantly differ from their estimated noise ceilings (p-values evaluated without corrections for multiple comparisons). In M1, only the measured correlations for the lowest and third highest frequency (Fig. 4d, pair 1 vs. 3) deviated significantly from the expected correlations (*p* = 0.035). In S1, the only significant deviations were for frequency condition pair 2 vs. 4 (Fig. 4h, *p* = 0.004). Although these deviations in M1 and S1 are statistically significant, the magnitude of the deviations were minor (average deviations in M1 and S1 = − 0.027). More importantly, the correlations between the RDMs measured at the lowest and highest activity level were not significantly different from the noise ceiling estimates. Therefore, these minor distortions do not appear to be systematic. These results demonstrate that the relationship between crossnobis dissimilarities remains relatively stable across a broad range of overall activity in sensorimotor and primary and secondary visual cortices.

**Figure 4:**
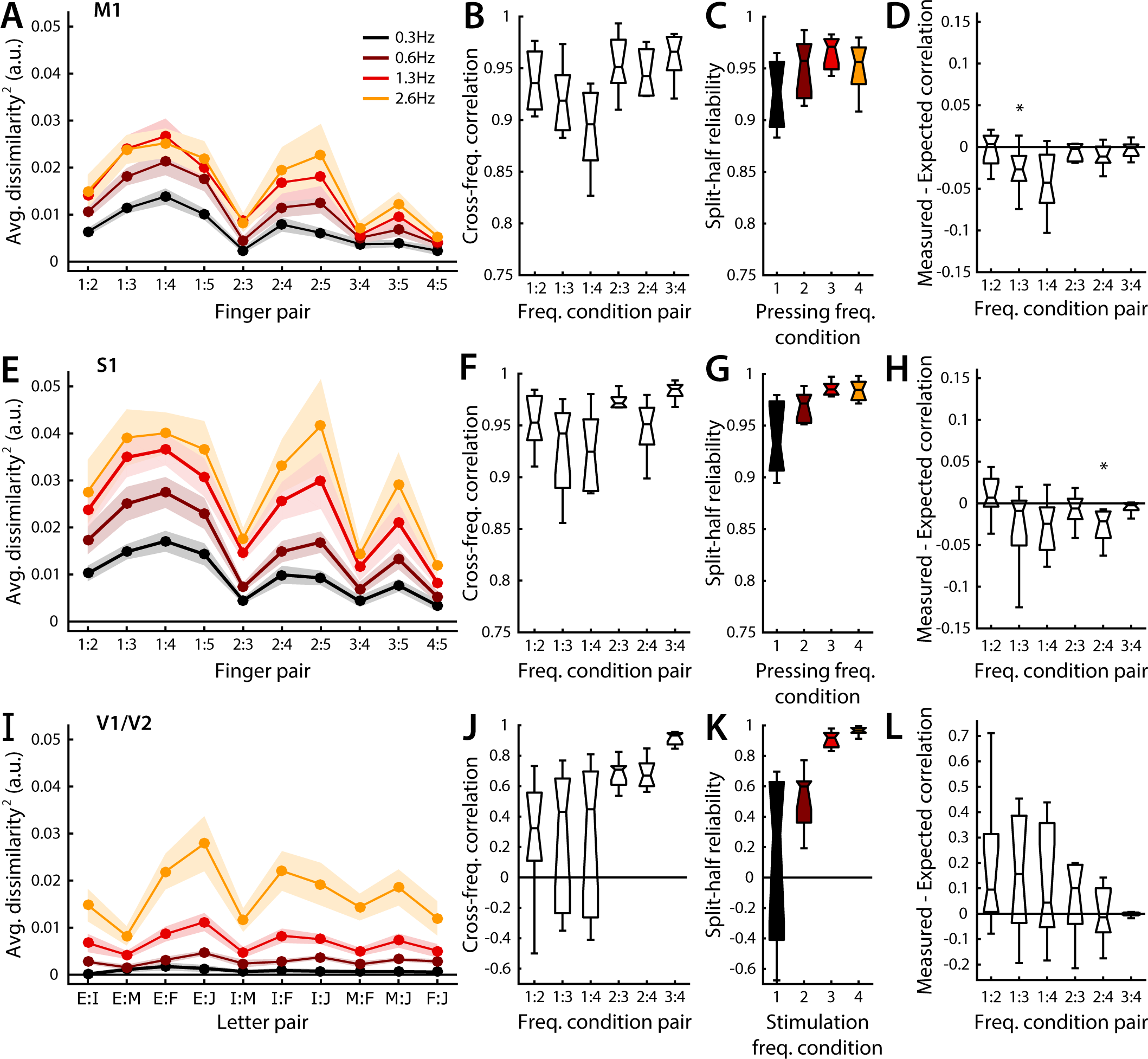
Stability of representational geometry across stimulation frequencies. (**A**) Average dissimilarity between all possible pairs of the 5 activity patterns measured at each frequency in M1. Colors indicate pressing/stimulation frequency, and shaded regions reflect s.e.m. (**B**) Cross-frequency correlations between dissimilarities in **A**. (**C**) Within participant split-half reliabilities (Pearson correlation with a forced intercept) of the dissimilarities depicted in **A**. (**D**) Differences between the expected correlation (noise ceiling, see section 2.9) and the measured cross-frequency correlations in (**B**). Negative values indicate the measured cross-frequency correlation (stability) is lower than what would be expected given the internal reliability of each RDM. Deviations from zero were evaluated with one-tailed signed rank tests. Asterisks indicate significant deviations (*p <* 0.05). (**E - H**) Results for S1. (**I - L**) Results for V1/V2. Note that the representational geometry in visual regions is different from the one found in M1/S1, reflecting the finger-to-letter assignment. Due to low stimulation intensity, visual regions have low reliabilities for low stimulation frequencies.

## 4. Discussion

Here we critically investigated whether possible non-linearities between neural activity and BOLD signal would lead to measurable distortions of the representational geometry as overall activity increases. We tested this assumption by stimulating both sensory-motor and visual regions at increasing frequencies. We assume that on the neural level, each repeated event should elicit approximately the same activity pattern. Therefore, the temporally integrated neural activity patterns should scale in an orderly fashion across frequencies. Importantly, this does not imply that neural activity would increase linearly with the number of events. Indeed, previous findings show that the neural response to subsequent finger taps is strongly attenuated in M1 (Hermes et al., 2012). This argues that the observed non-linear response in our data (Fig. 2b) can be attributed to a behavioral-to-neural rather than to a neural-to-BOLD non-linearity.

We found that an orderly scaling across frequencies could be observed in the BOLD activity patterns, even though the local activity increased over a large dynamic range – likely close to the achievable maximum for this paradigm. These findings corroborate previous studies that report linear coupling between neural activity and BOLD responses in M1/S1 (Siero et al., 2013), and V1 (Boynton et al., 1996; Heeger et al., 2000). However, our paper provides an important extension to these findings. Showing that BOLD signal and neural activity are linearly coupled on average in a region does not guarantee that the fine-grained activity patterns would also retain their representational geometry. For this, every individual voxel would have to exhibit linear scaling. By demonstrating the stability of the representational dissimilarities, we provide indirect evidence for this behavior to be true on the population level. Therefore, our findings present the first rigorous test that the commonly-held assumption of linear neural-to-BOLD coupling holds true at the level of multivariate activity patterns.

These findings have important implications for multivariate fMRI analyses. In general, most papers rely explicitly or implicitly on the assumption that the representational geometry as measured with fMRI veridically reflects the representational geometry of the underlying neural population code. While this strong assumption may still be violated for other reasons (see below), we show here that the representational geometry can be meaningfully compared across a relatively large range of overall activation levels. Thus, under appropriate conditions, representational geometries can be compared across different regions, individuals, or patient populations. Furthermore, from our data it appears justified to compare RDMs between different attentional states or different levels of learning, even if these differ strongly in their average activity.

For RSA, these findings also have implications for the choice of a dissimilarity measure. Not all measures make equally strong assumptions about the relation between the underlying neural representational geometry, and the one measured with fMRI. For example, a common practice in RSA is to evaluate rank-correlation between measured and predicted RDMs. Arguably, this approach makes inferences more robust against minor distortions in the measurement process (Kriegeskorte et al., 2008; Nili et al., 2014). Our results indicate that the exact ratio-relationship between dissimilarity measures can be meaningfully interpreted across a large range of average activation states. For this to be true, we of course need to use a dissimilarity measure that provides a meaningful zero point unbiased by noise – a condition met by the crossnobis estimator (Diedrichsen et al., 2016; Kriegeskorte and Diedrichsen, 2016; Walther et al., 2016). The additional information in the exact ratio-relationships allow for more powerful inferences about the underlying representations (Diedrichsen and Kriegeskorte, 2017).

The stability of the representational geometry is also good news for alternative approaches that test representational models. Pattern component modeling (PCM, Diedrichsen et al. 2011, 2017) and encoding models (Naselaris et al., 2011) make inferences about the underlying representational geometry in a very similar way to RSA (Diedrichsen and Kriegeskorte, 2017). Therefore, PCM and encoding approaches are subject to similar assumptions as RSA – and our findings generalize, such that inferences using these two methods will also be stable across activity levels.

Do these results suggest that one can make inferences about the representational geometry of the underlying neural population code from fMRI measures? While our results are reassuring in some aspects, there are two important caveats that we did not address in the current paper. First, fMRI samples neural activity with dramatic spatial averaging, even when using sub-millimeter resolution. Representations that exist at a finer spatial scale in the neural activity patterns will be under-represented in BOLD activity patterns, while representations at a large spatial scale will be over-represented (see Kriegeskorte and Diedrichsen 2016). Therefore, the representational geometry of BOLD activity patterns may differ systematically from the underlying neural code.

Secondly, the physiological processes underlying the BOLD signal and underlying extracellular neural recording are fundamentally different: While the BOLD signal reflects to a large degree the metabolically expensive processes of ion transport after excitatory postsynaptic potentials (Attwell and Iadecola, 2002; Attwell et al., 2010), extracellular recordings reflect neural spiking. Crudely spoken, therefore, BOLD reflects more the input to a region, while neural extracellular recordings reflect the output. Additionally, most extracellular recordings are biased towards large output neurons, as these provide the clearest extracellular signal (Firmin et al., 2014; Harris et al., 2016), whereas the BOLD signal indiscriminately averages metabolic activity. These important caveats need to be kept in mind when drawing parallels between representational analysis of extracellular recordings and BOLD signal.

While we were successful in driving the overall activity level in M1 and S1 across a large range, we did not achieve the same in the visual cortices. The activity and representations in V1/V2 at low stimulation frequency were unreliable. While be obtained substantially greater BOLD activity for the two highest frequencies, the activation levels were lower than those in M1 and S1. For these observed activity levels, however, we found good stability of the representational geometry.

### 4.1. Conclusion

One common assumption in multivariate fMRI analyses is that the relationship between activity patterns can be meaningfully interpreted. Possible non-linearities in neural-to-BOLD coupling, however, could lead to substantial distortions of multivariate measures when overall activity increases. Our results demonstrate that, across a broad range of overall activation states in M1, S1, and V1/V2, the ratio-relationships between pair-wise dissimilarities remain stable. This suggests that it is viable to leverage more powerful techniques, such as the use of cross-validated dissimilarities and likelihood-based RSA (Diedrichsen et al., 2016, 2017), for model comparison. The finding also applies to other multivariate techniques that analyze the relationships of BOLD activity patterns.

## Acknowledgements

The work was supported by a NSERC discovery grant (RGPIN-2016-04890) to JD. Functional imaging was supported partly by a Platform Support Grant from Brain Canada and the Canada First Research Excellence Fund (BrainsCAN). We thank Nikolaus Kriegeskorte, Timothy Verstynen, and Patrick Beukma for helpful comments on an earlier draft of the paper.

